# Sexual dimorphism in outcomes of non-muscle invasive bladder cancer: a role of CD163+ M2 macrophages, B cells and PD-L1 immune checkpoint

**DOI:** 10.1101/2021.01.23.427909

**Authors:** Stephen Chenard, Chelsea Jackson, Thiago Vidotto, Lina Chen, Céline Hardy, Tamara Jamaspishvilli, David Berman, D. Robert Siemens, Madhuri Koti

## Abstract

Non-muscle invasive bladder cancer (NMIBC) is significantly more common in men than women. However, female patients with NMIBC often present with more aggressive disease and do not respond as well to immunotherapy treatments. We hypothesized that sexual dimorphism in the tumor immune microenvironment (TIME) may contribute to the inferior clinical outcomes observed in female patients. To test this hypothesis, we interrogated the expression patterns of genes associated with specific immune cell types and immune regulatory pathways using tumor whole transcriptome profiles from male (n=357) and female (n=103) patients with NMIBC. High-grade tumors from female patients exhibited significantly increased expression of *CD40, CTLA4, PDCD1, LAG3* and *ICOS* immune checkpoint genes. Based on the significant differences in expression profiles of these genes and the cell types that most commonly express these in the TIME, we evaluated the density and spatial distribution of CD8+Ki67+ (activated cytotoxic T cells), FoxP3+ (regulatory T cells), CD103+ (tissue resident T cells), CD163+ (M2-like tumor associated macrophages), CD79a+ (B cells), PD-L1+ (Programmed-Death Ligand-1) and PD-1+ cells using multiplexed immunofluorescence in an independent cohort of 332 patient tumors on a tissue microarray (n=259 males and n=73 females). Tumors from female patients showed significantly higher infiltration of CD163+ macrophages and PD-L1+ cells compared to tumors from male patients. Notably, increased infiltration of CD163+ macrophages and CD79a+ B cells independently associated with decreased recurrence free survival. Not only do these results have the potential to inform the rational utilization of immunomodulatory therapies based on the TIME of both male and female patients with NMIBC, these novel findings highlight the necessity of considering sexual dimorphism in the design of future immunotherapy trials.

## Introduction

In 2018, bladder cancer was the 10th most common cancer worldwide, with an estimated 549,000 newly diagnosed cases and approximately 200,000 deaths caused by the disease [1]. Over 90% of bladder tumors arise from the urothelial cells that line the bladder. Localized urothelial carcinoma of the bladder (UCB) can be broadly categorized based on the depth of tumor invasion; approximately 75% of incident cases present as non-muscle-invasive bladder cancer (NMIBC), while the rest are classified as muscle-invasive disease [2].

The gold-standard first-line treatment for NMIBC is transurethral resection of bladder tumor (TURBT) surgery. Following surgical resection, patients diagnosed with high-risk features (such as high stage, grade, or tumor multifocality) [3] are best treated with intravesical Bacillus Calmette-Guérin (BCG) immunotherapy. BCG is a live-attenuated form of *Mycobacterium bovis (M. bovis)* and is commonly used as a vaccine for the prevention of tuberculosis. While NMIBC is four times more common in men, women suffer earlier recurrences following treatment with BCG immunotherapy and experience shorter progression-free survival (PFS) when compared to their male counterparts [4–6]. Clearly, patient sex (biological differences) and gender (social/behavioral differences) are associated with the incidence and clinical outcomes of NMIBC; however, these factors are understudied in biomarker and treatment design [7].

The pre-treatment tumor immune microenvironment (TIME) is a critical determinant of response to immunomodulation in several cancer types [8], and potentially plays a similar role in determining the response of bladder cancer patients to treatment with BCG immunotherapy [9,10]. The evolution of variable TIME states across the spectrum of non-inflamed (low/no immune infiltration) to inflamed (high immune cell infiltration) tumors depends on host and/or cancer cell intrinsic factors that influence the recruitment and functional states of immune cells [11]. Aligning with this concept and in the context of contemporary immunomodulatory agents including the PD-1/PD-L1 targeting immune checkpoint blockade therapy, patient biological sex has recently emerged as an important factor in response to treatment [12–14]. Under normal physiological conditions, macrophages are the most abundant resident innate immune cells within the bladder mucosa. Pre-clinical studies in murine models have shown that females exhibit higher magnitude of immune responses to urinary pathogens than males [15]. Advancing age, microbial challenges, host genetics and post-menopausal hormonal changes also contribute to increased recruitment of adaptive immune cell populations, within the bladders of females. Despite mounting evidence of the importance of these cell types in response to immunomodulatory therapy in several other cancers, their sex-specific roles and functional states within the NMIBC TIME have yet to be fully elucidated.

We hypothesized that sexual dimorphism in clinical outcomes of NMIBC are driven by differences in the TIME. To test this hypothesis, we evaluated the association between patient sex and available clinical outcomes in two independent cohorts of NMIBC; UROMOL (publicly available dataset of tumors from 460 patients) [16] and Kingston Health Sciences Center (KHSC; 332 patients). We then evaluated the tumor transcriptome profiles of the UROMOL cohort to determine sex associated differences in immune regulatory genes. Guided by the findings from immune regulatory gene expression patterns, we further investigated the density and spatial distribution of selected immune cell populations and immune checkpoint proteins in tumors from male and female patients in the KHSC cohort. Findings from our study provide the first evidence for sex associated differences in the TIME of NMIBC.

## Results

### Female patients with high grade NMIBC tumors exhibit shorter progression free survival

We evaluated the association between patient sex and available clinical outcomes in two independent cohorts of NMIBC; UROMOL (460 patients) [16] and KHSC (332 patients). Clinical features of the KHSC cohort are presented in **Table S1**, whereas those for the UROMOL cohort were previously reported [16]. In the KHSC cohort, 22% of the patients were female, 60% of the entire cohort had high-grade disease (Supplementary **Table S2**) at original presentation and the majority (88%) did not have evidence of BCG immunotherapy prior to collection of their specimens (BCG naïve). Approximately 38% of patients received one or more doses of BCG (**Table S1**). Clinical care of all patients was coordinated at a single treatment clinic and decisions regarding adjuvant therapies were based on AUA risk score. The UROMOL cohort had a similar proportion of female patients (22%). The proportions of patients who underwent BCG immunotherapy was 19% and 54% in the UROMOL and KHSC cohorts, respectively. In alignment with previous reports on sex associated differential outcomes, female patients with high-grade NMIBC had a shorter PFS compared to males in both the cohorts (**Figure 1A and 1B**).

**Figure 1.**
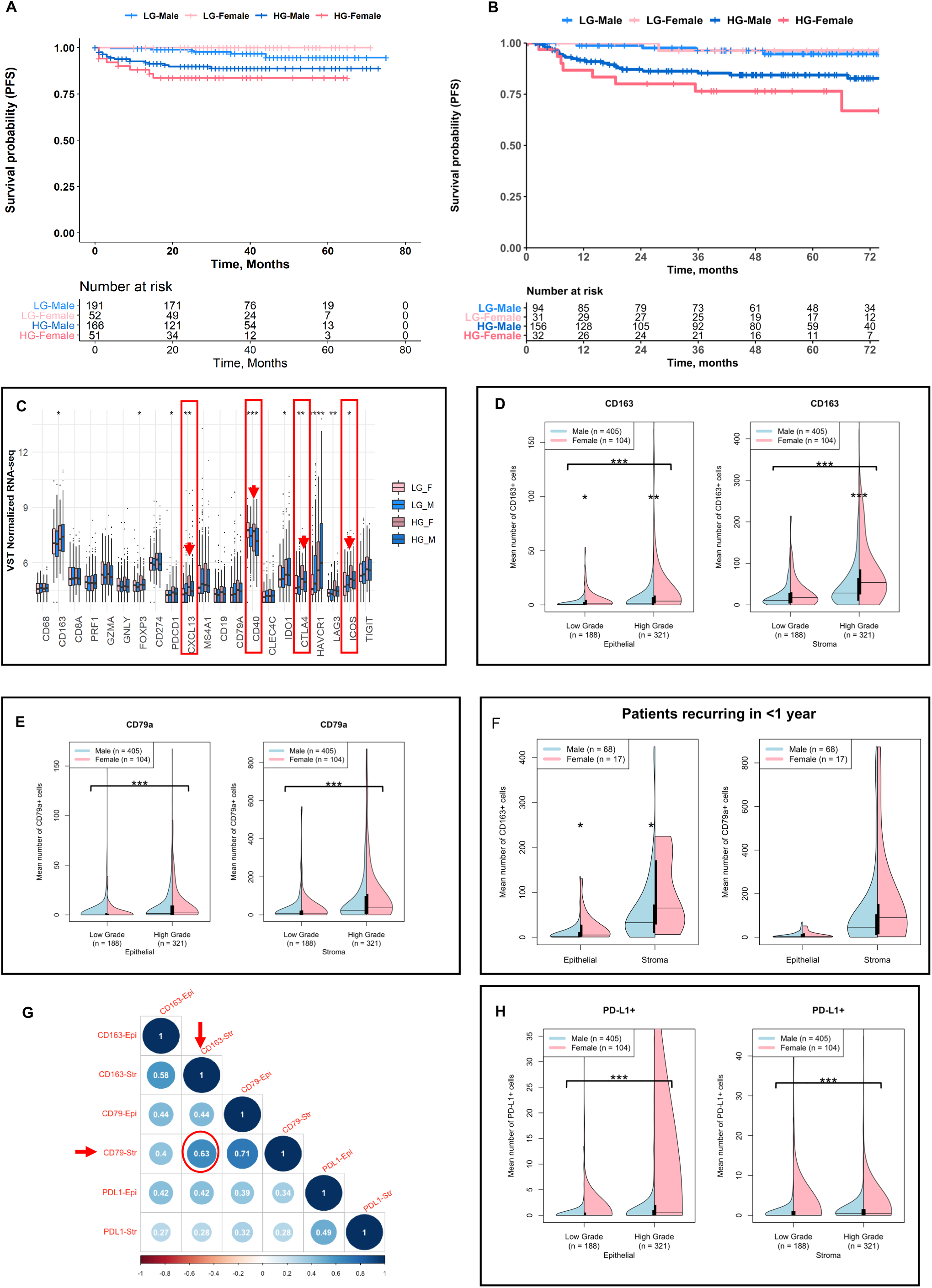
Patients with NMIBC exhibit sexual dimorphism in progression free survival and tumor immune microenvironment. (A-B) Kaplan Meier survival curves showing female patients with high-grade NMIBC experienced significantly (P<0.001) shorter PFS than their male counterparts and patients with low-grade tumors, in both the UROMOL (A) and KHSC (B) cohorts. Kaplan-Meier survival analyses was performed using log-rank statistics with survival and survminer packages. (C) High-grade tumors from female patients in the UROMOL cohort exhibit significantly increased expression of immunoregulatory genes, *CXCL13, PDCD1, CD40, CTLA4*, and *ICOS*. VST-normalized genes between four cohorts: high-grade cancer in females (HG_F), low-grade cancer in females (LG_F), high-grade cancer in males (HG_M), and low-grade cancer in males (LG_M) are shown. The Kruskal-Wallis test was employed to determine statistically significant differences. (D-E) Violin plots of mean cell counts for CD163+, CD79a+ cell and PD-L1+ cell populations respectively, in the epithelial (left) and stromal (right) compartments of tumors from the KHSC cohort. Plots stratified by low-grade and high-grade samples for males (blue) versus females (pink). Significant differences signified by asterisks between grades (with bracket) and between sexes (no bracket) as determined by Mann-Whitney-U test. ***p-value < 0.001, **p-value < 0.01, *p-value < 0.05. (F) In a subset of patients from the KHSC cohort that recurred in <1 year, female patients had significantly higher density of CD163+ cells in the stroma and epithelium, while there were no significant sex-associated differences in CD79a+ cells. *p-value < 0.05. (G) Spearman correlation plot for CD163+, CD79a+ and PD-L1+ populations in the epithelial (Epi) and stromal (Str) compartments of tumors from the KHSC cohort. Dark blue indicates a positive correlation coefficient (>1), dark red indicates a negative correlation coefficient (<1). (H) Violin plots of mean cell counts for PD-L1+ cell populations in the epithelial (left) and stromal (right) compartments of tumors from the KHSC cohort. Plots stratified by low-grade and high-grade samples for males (blue) versus females (pink). Differences between grades in each compartment as determined by Mann-Whitney-U test ***p-value < 0.001.

### High grade tumors from female patients with NMIBC exhibit increased expression of immune regulatory genes

Using whole transcriptome profiles of tumors in the UROMOL cohort (n= 103 female and 357 male), a targeted analysis was performed to determine grade- and sex-associated differential expression patterns of genes identifying immune cell phenotypes (specifically macrophages, T cells, and B cells), their functional states, and immune-regulatory functions [17]. Significantly higher expression of the immune checkpoint genes *CTLA4, PDCD1, LAG3* and *ICOS*, were observed in high-grade tumors from females compared to those from males and compared to low-grade tumors from both sexes (**Figure 1C**). Importantly, transcript levels of B cell recruiting chemokine, *CXC ligand 13* (*CXCL13*) and B cell surface-associated molecule, *CD40* were significantly increased in high-grade tumors from female patients (**Figure 1C**).

### Tumors from female patients with NMIBC have increased CD163+ M2-like macrophages, CD79a+ B cells and higher PD-L1 immune checkpoint protein expression

Based on the differences in immunoregulatory gene expression profiles investigated in the UROMOL cohort, we next evaluated sex-associated differences in the density and localization of a subset of immune cells that are known to express the relevant phenotypic markers and immune checkpoint proteins, and have the potential to secrete the cytokine CXCL13. As such, we investigated the epithelial and stromal density profiles of T helper (CD3+CD8-), T cytotoxic (CD8+Ki67-, activated CD8+Ki67+), T regulatory (CD3+CD8-FOXP3+), tissue resident T (CD103+), B-cells (CD79a), M2-like tumor associated macrophages (TAMs, CD163+) and immune checkpoint proteins (PD-L1 and PD-1) in tumors from the KHSC cohort.

Interestingly, among all the immune cell phenotypes, only CD163+ M2-like TAMs demonstrated statistically significant differences in the density and localization between both low grade and high-grade tumors from male and female patients. A significantly higher infiltration of CD163+ TAMs was seen in the epithelial compartment of low-grade tumors (p=0.012) and in both epithelial and stromal compartments of high-grade tumors (p= 0.001, p<0.001, respectively) from female patients compared to those from male patients (**Figure 1D**). Density of CD79a+ B cells was not significantly different between sexes; however, CD79a+ B cell infiltration was higher in the epithelial and stromal compartments (p<0.001) of high-grade tumors compared to low-grade tumors from both sexes (**Figure 1E**). Importantly, tumors from female patients exhibiting recurrence within one year showed significantly higher density of CD163+ TAMs compared to those from males (**Figure 1F**). With respect to CD79a+ B cells, a trend towards higher infiltration was observed in tumors from female patients who recurred within 1 year of BCG treatment (**Figure 1F**). Spearman correlation analysis showed a moderate positive correlation (r=0.63) between stromal CD79a+ B cells and CD163+ TAMs (**Figure 1G**).

PD-L1 protein expression was significantly higher in both the epithelial and stromal compartments of high-grade tumors compared to low-grade tumors (**Figure 1H**; p<0.001). However, apparent sex-associated differences in PD-L1 protein expression (especially in the epithelial compartment) were likely not statistically significant due to the wide variability in the proportion of PD-L1+ cells.

A subset analysis of cases was performed on patients that had no evidence of previous intravesical BCG immunotherapy (BCG naïve) by excluding from the entire cohort those patients that had documented BCG prior to specimen collection or where that data could not be definitely determined. Trends in infiltration profiles of CD163 and CD79a between grades and between sexes were consistent with those seen in the whole cohort (**Supplementary Figures 2A-C**) although in this analysis expression of PD-L1 protein in the epithelial compartment of low-grade tumors from females was significantly higher compared to males (p=0.04).

### T helper and regulatory cells exhibit significant sex differences in density and spatial distribution within the NMIBC TIME

While we observed no sex-associated differences in the density and spatial organization of CD8+ cytotoxic T cells, CD3+CD8-T helper cells and CD3+CD8-FoxP3+ T regulatory cells were significantly higher in low-grade tumors from female patients compared to those from males (Supplementary Fig. 3A and B). Such differences were not observed in high-grade tumors.

### Increased density of CD163+ M2-like macrophages and CD79a+ B cells associates with early recurrence in NMIBC

Kaplan-Meier analysis for all patients with high-grade NMIBC (n=170) in the KHSC cohort showed that, irrespective of their localization in the stroma or epithelium, higher density (Supplementary **Table S2**) of CD163+ M2-like TAMs (**Figure 2A**) and CD79a+ B cells (**Figures 2B**) independently associated with shorter recurrence free survival (RFS). This association was observed in both male and female patients supporting the notion that rather than being a sexually dimorphic epiphenomenon, these cells may play a functional role in tumor recurrence (**Figures 2C and 2D**). Notably, these differences in RFS remained consistent using similar thresholds for both CD79a+ and CD163+ cells, in all patients with high-grade tumors (**Supplementary Figures 4a, 4b, 5a and 5b**).

**Figure 2.**
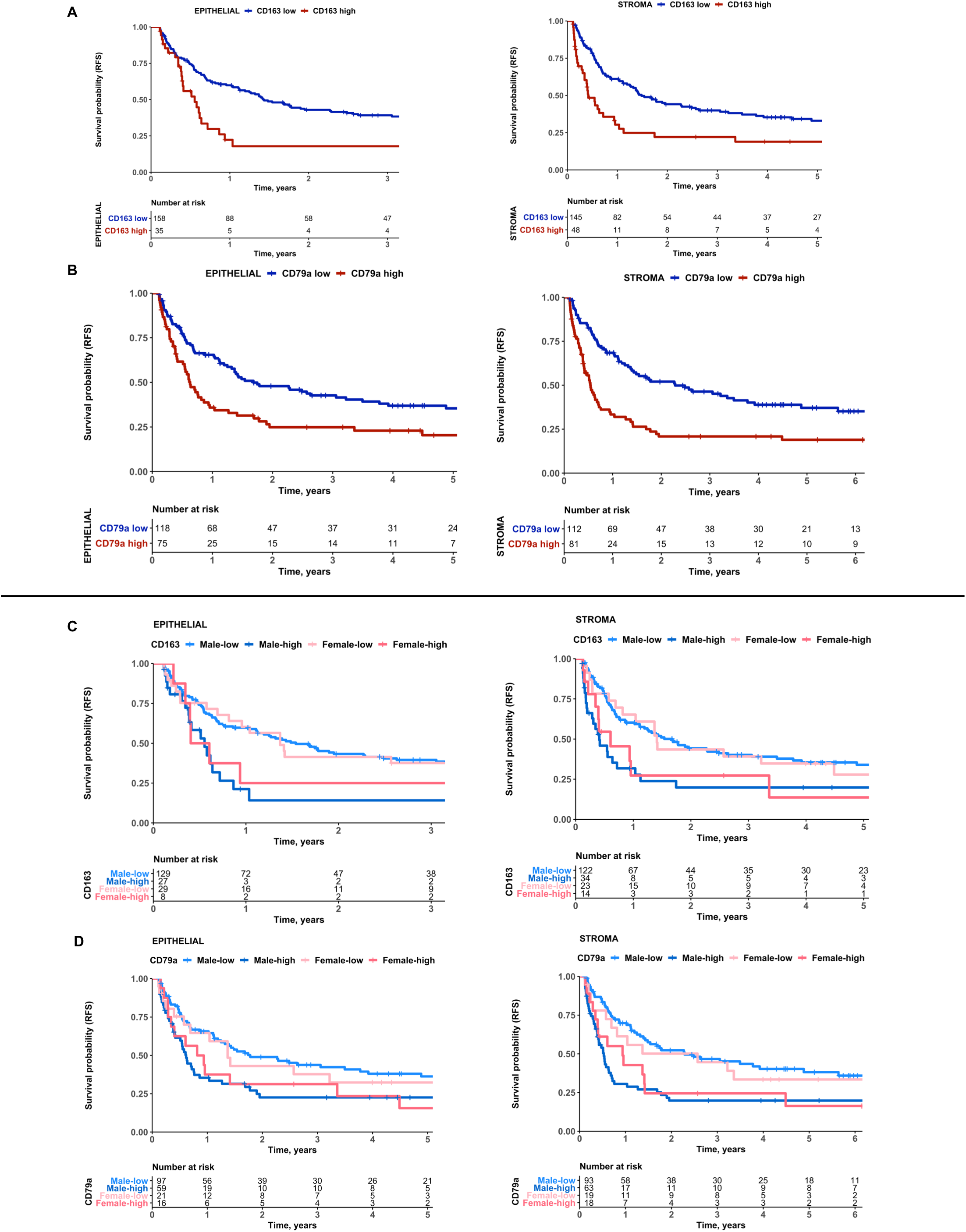
Recurrence free survival (RFS) in high-grade non-muscle invasive bladder cancer is associated with patient sex and tumor immune microenvironment. (A) Kaplan Meier survival curves for RFS of patients with high-grade NMIBC (n=193) based on log-rank optimized thresholds for density of CD163+ cells in tumor epithelial (left; P<0.001) and stromal (right; P<0.001) compartments. Of this entire cohort, 74% had evidence of adequate BCG therapy after specimen collection (TURBT). (B) Kaplan Meier survival curves showing RFS of high-grade patients based on log-rank optimized thresholds for epithelial (left; P<0.01) and stromal (right; P<0.0001) CD79a+ cells. (C) Kaplan Meier survival curves showing RFS of high-grade patients based on log-rank optimized thresholds for stromal (left; P<0.01) and epithelial (right; P=0.014) CD163+ cells stratified by sex. High vs low stromal CD163+ cells defined as > 64 or < 64 cells, respectively. High versus low epithelial CD163+ cells defined as > 12 or < 12 cells, respectively. (D) Kaplan Meier survival curves showing RFS of high-grade patients based on log-rank optimized thresholds for stromal (left; P<0.001) and epithelial (right; P=0.028) CD79a+ cells stratified by sex. High versus low stromal CD79a+ cells defined as > 35 or < 35 cells, respectively. High vs low epithelial CD79a+ cells defined as >3 or < 3 cells, respectively.

## Discussion

In the current study, we report the first demonstration of sexual dimorphism in the TIME of NMIBC. In concordance with previous reports [4–6], we found that female patients with high-grade NMIBC suffer from shorter progression free survival as compared to their male counterparts. Analysis of a panel of immune regulatory genes (both stimulatory and inhibitory) in tumors from the UROMOL cohort demonstrated increased expression of immune checkpoint genes, and those associated with B cell recruitment and function in high-grade tumors from females compared to those from males. These alterations are indicative of an exhausted immune landscape following an increased activation within the NMIBC TIME. Given the dynamic nature of immune checkpoint gene expression, variability in checkpoint co-expression, and an expected lack of direct correlation with protein level expression, we investigated the profiles of cell types known to express these molecules following activation or exhaustion.

Our finding of higher infiltration of CD163+ M2-like suppressive TAMs in tumors from female patients may partially explain the overall poor clinical outcomes experienced by female patients following BCG immunotherapy. A significantly increased density of M2-like TAMs in tumors from female patients may be driven by differences dictated by sexual dimorphism in overall bladder mucosal immune physiology [15]. Indeed, higher infiltration of tumors by CD163+ M2-like TAMs has been previously reported to be associated with poor clinical outcomes in NMIBC [18]. It is also known that bladder cancer cells induce the polarization of tissue-resident and reactive macrophages [19], potentially influencing tumor progression and treatment response [20]. Our novel findings of the higher density of B cells in tumors from female patients are reflective of the established physiological links between M2-like TAMs and B cells [21]; co-stimulation of M2-like macrophages with bacterial lipopolysaccharide and IL-10 induces production of CXCL13, a chemokine critical for B cell recruitment [22,23]. Given the increased incidence of urinary tract infections in women – and the cancer cell-induced polarization of TAMs towards an M2-like phenotype [19] – it is plausible that increased engagement of M2-like TAMs by urinary pathogens leads to increased CXCL13 secretion and B cell recruitment in tumors of female NMIBC patients. Eventually, these recruited cells acquire a dysfunctional or exhausted phenotype, persisting in an immunological stalemate within the NMIBC TIME. This observation is further strengthened by our *in silico* transcriptomic analysis showing increased expression of *CXCL13*, however, further mechanistic evidence is warranted. Similarly, it is also known that IL-10 secreted by B regulatory cells inhibits macrophage activation and polarizes them towards an M2-like phenotype [24]. Indeed, under normal physiological conditions, females exhibit higher proportions of B cells and increased responsiveness to BCG vaccination [25,26]. However, given the older age of patients with NMIBC, it is possible that a reduced naïve B cell pool in the periphery [27] compromises the desired responses in the NMIBC scenario following local BCG administration in contrast to responses associated with infant vaccination. We acknowledge that the static TIME states evaluated in this study, however, do not provide definitive evidence supporting these temporal phenomena and warrant further investigation.

Another novel finding from our study is the inverse association of B cells and CD163+ M2-like TAMs with recurrence free survival in BCG naïve patients. This finding has significant implications in advancing the current state of knowledge on the antigen presenting function of these cells following encounter with BCG bacteria upon their intra-vesical administration. For example, it is known that M2-like macrophages exhibit tolerance and are unable to secrete CXCL10 [28], which limits their ability to recruit immune cells to the TIME. However, BCG treatment has been shown to re-program M2 macrophages [29] that may lead to some degree of the observed anti-tumor responses. B cells have been shown to be indicators of good prognosis in certain cancers [30,31]. However, their anti-tumor roles likely depend either on their antibody producing or antigen presentation function, neither of which are well understood in NMIBC. The negative prognostic association of B cells, as observed in this study, indicates their potentially exhausted or regulatory phenotype. A comprehensive analysis of their functional states is needed to define their tumor killing or promoting roles.

Another important finding from this study is the sex-associated difference in PD-L1 protein expression. PD-L1 is known to be expressed on a wide variety of immune cells and cancer cells and is partially regulated by estrogen and X-linked microRNAs [32,33]. Indeed, pre-treatment PD-L1 expression was recently shown to be predictive of response to BCG [34]. The current finding suggesting overall higher PD-L1 expression in tumors from females may inform more precise use of combination therapies targeting this immune checkpoint in female patients despite the challenges due to the dynamic nature of its expression.

The current findings should inform immunotherapy trials that are adequately powered to evaluate responses in female patients with NMIBC. Increased abundance of CD163+ M2-like macrophages may also explain the compounding effects of these suppressive factors that impart an aggressive behavior leading to poor clinical outcomes experienced by female patients relative to males. Several trials targeting CD40 or CSF1R (M2 TAM targeting) in combination with PD-L1 immune checkpoint blockade are underway in a variety of cancers [35], outcomes of which might inform their potential use in NMIBC.

Our study is not without limitations. PD-L1 immune checkpoint is expressed on a wide variety of cells including cancer and immune cells in the TIME. Future studies evaluating the co-localization of immune checkpoint should derive definitive information on cell types expressing this immune checkpoint. Furthermore, a more comprehensive evaluation of the functional states of immune cells and immunogenomic correlates driving their recruitment via cell intrinsic interferon activation, is needed to guide more precise therapeutic targeting. Nevertheless, findings from this study have significant implications in ongoing immune checkpoint blockade trials where sex-associated pre-treatment tumor immune landscape could inform their precise use in drug sequencing.

## Methods

### Patient Cohorts and Clinical Data

This study was approved by the Ethics Review Board at Queen’s University. A cohort of 509 NMIBC archival transurethral bladder tumors (TURBTs) from 332 patients between 2008-2016 was collected from Kingston Health Sciences Center (KHSC). Stage Ta and T1 tumors were included following pathological review (DMB and LC), using the WHO 2016 grading system [36]. Clinical details of the UROMOL cohort were previously reported by Hedegaard et al. 2016 [16]. Six tissue microarrays (TMA) were constructed with duplicate 1.0 mm cores. Recurrence was defined as time from each patient’s earliest TURBT resection to next malignant diagnosis. Operative notes were reviewed to exclude re-resections as recurrences.

### Multiplex Immunofluorescence Staining for Immune Markers

TMAs were stained with three panels of primary-conjugated fluorescent antibodies. The first panel contained antibodies against CD3+, CD8+, Ki67+, and FoxP3+ cells. The second panel contained antibodies against PD-1+, PD-L1+, and CK5+ cells. The third panel contained antibodies against CD163+, CD79a+, CD103+, and GATA3+ cells. Expression for the following cell types and immune checkpoints were evaluated in the epithelial and stromal compartments: T-cells (CD3+CD8+Ki67, CD3+CD8+Ki67+, CD3+CD8-FOXP3-, CD3+CD8-FOXP3+), immune checkpoint (PD-1+, PDL1+, CK5+PDL1+), B-cells (CD79a+), M2-like TAMs (CD163+), and tissue-resident T cells (CD103+). The presence of these immune markers was evaluated for each core by automated multiplex immunofluorescent staining at the Molecular and Cellular Immunology Core (MCIC) facility, BC Cancer Agency. All antibodies were provided by Biocare Medical (Pacheco, CA, USA) and distributed by Inter Medico (Markham, ON, CAN).

### Automated Scoring of Multiplex Immunofluorescence Staining for Immune Markers

The stained TMA sections were scanned using the Vectra multispectral imaging system. Tissue segmentation into stromal and epithelial compartments was performed using PerkinElmer’s InForm software. Ten randomly selected cores were utilized to train three independent algorithms in PerkinElmer’s InForm software package. The InForm software identified positive pixels for all of the selected immune markers using each of the three independent algorithms. The average of the three independent algorithms was taken for all three sets of immune markers in each core. Cores that were missing ≥ 75% of the tissue were excluded from further analyses.

### Manual Validation of Automated Scoring of Multiplex Immunofluorescence Staining

Standard deviation between the three algorithms was calculated for each of the immune markers within both the epithelial and stromal compartments. Outliers were identified and cross-referenced with the composite image of the corresponding TMA core to verify the discrepancy before excluding the outlying data points from further analysis. Any quantifications that included misidentification of histological artifacts as immune cells by any of the three algorithms were also excluded from analysis. Algorithms that consistently over-called or under-called the immune marker quantification were excluded from analysis. Further visual validation of the automated scoring was performed for randomly selected TMA cores.

### RNAseq Gene Expression Analysis

Raw RNAseq data from 460 NMIBC samples [16], were downloaded (https://www.ebi.ac.uk/ega/studies/EGAS00001001236), and VST normalized data were obtained by employing the vst function on DESeq2 in R v4.0.1. Further, we compiled a list comprising immune-cell markers and regulatory genes based on our previous report [17]. We then compared the expression of these VST-normalized genes between four groups of patients: high-grade females, low-grade females, high-grade males, and low-grade males. The Kruskal-Wallis test was employed to determine significant differences between the two four groups.

### Statistical Analysis

Analyses were conducted using R version 3.5.3. Kaplan-Meier curves were plotted using log-rank statistics with survival and survminer packages. Follow-up time for Kaplan-Meier curves ended when 10% of patients remained in each group [37]. Log-rank statistics were used to optimize thresholds for ideal number of CD163+ M2-like TAMs/CD79a+ B cells associated with shorter recurrence-free survival, using the maxstat package in R. Optimal thresholds are summarized in **Supplemental Table S3**.

## Acknowledgements

This work is supported by the Early Researcher Award; Ontario Ministry of Research Innovation and Science and the Mary and Mihran Basmajian award for Excellence in Health Research,, Queen’s University to MK and SEAMO Innovation award to DRS and MK.

## Supplementary Figures

**Supplementary Figure 1. Multiplex immunofluorescence staining and automated tissue segmentation of representative tumor core**. A composite view of a representative tumor core, highlighting antibody panels distinguishing CD3+CD8+Ki67+/- and CD3+CD8-FoxP3-cells (A); CD79a+ B, CD103+ T resident and CD163+ M2-like TAMs (B); PD-1+, PD-L1+ and CK5+ cells (C). PerkinElmer’s Inform software based automated segmentation of tumor core into epithelial (red) and stromal (green) compartments prior to automated scoring of positively stained cells (D).

**Supplementary Figure 2. Profiles of CD163, CD79a and PD-L1 in tumors from BCG naïve patients**

Violin plots of mean cell counts for CD163+ (A), CD79a+ (B) and PD-L1+ cell (C) populations respectively, in the epithelial (left) and stromal (right) compartments of tumors from the KHSC cohort with no evidence of BCG immunotherapy prior to collection of their specimens (BCG naïve). Asterisks indicate level of significance as determined by Mann-Whitney-U statistics: ***p-value < 0.001 **p-value < 0.01 *p-value < 0.05.

**Supplementary Figure 3. Profiles of T helper (3A) and T regulatory (3B) cells in tumors from BCG naïve patients and overall cohort**

Violin plots of mean cell counts for CD3+CD8-T helper (A) and CD3+CD8-FoxP3+ T regulatory cells (B) in BCG naïve and overall cohort. Plots stratified by low-grade and high-grade samples for males (blue) versus females (pink). Asterisks indicate level of significance as determined by Mann-Whitney-U statistics: ***p-value < 0.001 **p-value < 0.01 *p-value < 0.05.

**Supplementary Figure 4. CD79a+ B cell density is associated with recurrence free survival in patients with high-grade NMIBC and no prior history of BCG before specimen collection (BCG naïve)**.

(a) Recurrence-free survival of a subset of patients with high-grade disease and no previous BCG therapy prior to specimen collection (n=170). Within this cohort 74% had evidence of adequate BCG therapy after specimen collection (TURBT). Based on log-rank optimized cut-offs for stromal (left; p<0.001) and epithelial (right; p<0.056) CD79a+ cells stratified by males (blue) versus females (pink).

(b) Recurrence-free survival based on log-rank optimized cut-offs for stromal (left) and epithelial (right) CD79a+ cells stratified by high CD79a+ cells versus low CD79a+ cells. High versus low stromal CD79a+ cells defined as > 35 or < 35 cells, respectively. High vs low epithelial CD79a+ cells defined as > 3 or < 3 cells, respectively. Associated p-values for the epithelial and stromal compartments is <0.01 and <0.0001, respectively.

**Supplementary Figure 4. Higher CD163+ cell infiltration is associated with shorter recurrence free survival in BCG-naïve patients with high-grade NMIBC**.

(a) Recurrence-free survival of high-grade, BCG naïve patients (n=170) based on log-rank optimized cut-offs for stromal (left; P=0.017) and epithelial (right; 0.057) CD7163+ cells stratified by males (blue) versus females (pink).

(b) Recurrence-free survival based on log-rank optimized cut-offs for stromal (left) and epithelial (right) CD163+ cells stratified by high CD163+ cells versus low CD163+ cells. High versus low stromal CD163+ cells defined as > 64 or < 64 cells, respectively. High vs low epithelial CD163+ cells defined as > 12 or < 12 cells, respectively. Associated p-values for the epithelial (right) and stromal (left) compartments are both p<0.01.

## Supplementary Tables

**S1. Clinical characteristics of patients in the KHSC cohort (n=332)**.

**S2. Sample characteristics (grade, stage and sex) for overall KHSC cohort (n = 509)**

**S3. Optimized log-rank thresholds for recurrence-free survival for individual immune markers**

## References

[1] Bray F, Ferlay J, Soerjomataram I, Siegel RL, Torre LA, Jemal A. Global cancer statistics 2018: GLOBOCAN estimates of incidence and mortality worldwide for 36 cancers in 185 countries. CA Cancer J Clin 2018;68:394–424. https://doi.org/10.3322/caac.21492.

[2] Woldu SL, Bagrodia A, Lotan Y. Guideline of guidelines: non-muscle-invasive bladder cancer. BJU Int 2017;119:371–80. https://doi.org/10.1111/bju.13760.

[3] Sylvester RJ, van der Meijden APM, Oosterlinck W, Witjes JA, Bouffioux C, Denis L, et al. Predicting recurrence and progression in individual patients with stage Ta T1 bladder cancer using EORTC risk tables: a combined analysis of 2596 patients from seven EORTC trials. Eur Urol 2006;49:466–7. https://doi.org/10.1016/j.eururo.2005.12.031.

[4] Uhlig A, Strauss A, Seif Amir Hosseini A, Lotz J, Trojan L, Schmid M, et al. Gender-specific Differences in Recurrence of Non-muscle-invasive Bladder Cancer: A Systematic Review and Meta-analysis. Eur Urol Focus 2018;4:924–36. https://doi.org/10.1016/j.euf.2017.08.007.

[5] Marks P, Soave A, Shariat SF, Fajkovic H, Fisch M, Rink M. Female with bladder cancer: what and why is there a difference? Transl Androl Urol 2016;5:668–82. https://doi.org/10.21037/tau.2016.03.22.

[6] Kluth LA, Fajkovic H, Xylinas E, Crivelli JJ, Passoni N, Rouprêt M, et al. Female gender is associated with higher risk of disease recurrence in patients with primary T1 high-grade urothelial carcinoma of the bladder. World J Urol 2013;31:1029–36. https://doi.org/10.1007/s00345-012-0996-9.

[7] Saginala K, Barsouk A, Aluru JS, Rawla P, Padala SA, Barsouk A. Epidemiology of Bladder Cancer. Med Sci (Basel, Switzerland) 2020;8. https://doi.org/10.3390/medsci8010015.

[8] Binnewies M, Roberts EW, Kersten K, Chan V, Fearon DF, Merad M, et al. Understanding the tumor immune microenvironment (TIME) for effective therapy. Nat Med 2018;24:541–50. https://doi.org/10.1038/s41591-018-0014-x.

[9] Annels NE, Simpson GR, Pandha H. Modifying the Non-muscle Invasive Bladder Cancer Immune Microenvironment for Optimal Therapeutic Response. Front Oncol 2020;10:175. https://doi.org/10.3389/fonc.2020.00175.

[10] Roumiguié M, Compérat E, Chaltiel L, Nouhaud FX, Verhoest G, Masson-Lecomte A, et al. PD-L1 expression and pattern of immune cells in pre-treatment specimens are associated with disease-free survival for HR-NMIBC undergoing BCG treatment. World J Urol 2020. https://doi.org/10.1007/s00345-020-03329-2.

[11] Giraldo NA, Sanchez-Salas R, Peske JD, Vano Y, Becht E, Petitprez F, et al. The clinical role of the TME in solid cancer. Br J Cancer 2019;120:45–53. https://doi.org/10.1038/s41416-018-0327-z.

[12] Conforti F, Pala L, Bagnardi V, Pas T De, Martinetti M, Viale G, et al. Articles Cancer immunotherapy efficacy and patients’ sex?: a systematic review and meta-analysis 2018;4:1–10. https://doi.org/10.1016/S1470-2045(18)30261-4.

[13] Wang C, Qiao W, Jiang Y, Zhu M, Shao J, Ren P, et al. Effect of sex on the efficacy of patients receiving immune checkpoint inhibitors in advanced non-small cell lung cancer. Cancer Med 2019;8:4023–31. https://doi.org/10.1002/cam4.2280.

[14] Wu Y, Ju Q, Jia K, Yu J, Shi H, Wu H, et al. Correlation between sex and efficacy of immune checkpoint inhibitors (PD-1 and CTLA-4 inhibitors). Int J Cancer 2018;143:45– 51. https://doi.org/10.1002/ijc.31301.

[15] Zychlinsky Scharff A, Rousseau M, Lacerda Mariano L, Canton T, Consiglio CR, Albert ML, et al. Sex differences in IL-17 contribute to chronicity in male versus female urinary tract infection. JCI Insight 2019;5. https://doi.org/10.1172/jci.insight.122998.

[16] Hedegaard J, Lamy P, Nordentoft I, Algaba F, Høyer S, Ulhøi BP, et al. Comprehensive Transcriptional Analysis of Early-Stage Urothelial Carcinoma. Cancer Cell 2016;30:27– 42. https://doi.org/10.1016/J.CCELL.2016.05.004.

[17] Vidotto T, Nersesian S, Graham C, Siemens DR, Koti M. DNA damage repair gene mutations and their association with tumor immune regulatory gene expression in muscle invasive bladder cancer subtypes. J Immunother Cancer 2019;7:148. https://doi.org/10.1186/s40425-019-0619-8.

[18] Wu S-Q, Xu R, Li X-F, Zhao X-K, Qian B-Z. Prognostic roles of tumor associated macrophages in bladder cancer: a system review and meta-analysis. Oncotarget 2018;9:25294–303. https://doi.org/10.18632/oncotarget.25334.

[19] Martínez VG, Rubio C, Martínez-Fernández M, Segovia C, López-Calderón F, Garín MI, et al. BMP4 Induces M2 Macrophage Polarization and Favors Tumor Progression in Bladder Cancer. Clin Cancer Res an Off J Am Assoc Cancer Res 2017;23:7388–99. https://doi.org/10.1158/1078-0432.CCR-17-1004.

[20] Suriano F, Santini D, Perrone G, Amato M, Vincenzi B, Tonini G, et al. Tumor associated macrophages polarization dictates the efficacy of BCG instillation in non-muscle invasive urothelial bladder cancer. J Exp Clin Cancer Res 2013;32:87. https://doi.org/10.1186/1756-9966-32-87.

[21] Mantovani A. B cells and macrophages in cancer: yin and yang. Nat Med 2011;17:285–6. https://doi.org/10.1038/nm0311-285.

[22] Vidyarthi A, Agnihotri T, Khan N, Singh S, Tewari MK, Radotra BD, et al. Predominance of M2 macrophages in gliomas leads to the suppression of local and systemic immunity. Cancer Immunol Immunother 2019;68:1995–2004. https://doi.org/10.1007/s00262-019-02423-8.

[23] Mantovani A, Sica A, Sozzani S, Allavena P, Vecchi A, Locati M. The chemokine system in diverse forms of macrophage activation and polarization. Trends Immunol 2004;25:677–86. https://doi.org/10.1016/j.it.2004.09.015.

[24] Fehres CM, van Uden NO, Yeremenko NG, Fernandez L, Franco Salinas G, van Duivenvoorde LM, et al. APRIL Induces a Novel Subset of IgA(+) Regulatory B Cells That Suppress Inflammation via Expression of IL-10 and PD-L1. Front Immunol 2019;10:1368. https://doi.org/10.3389/fimmu.2019.01368.

[25] Birk NM, Nissen TN, Kjærgaard J, Hartling HJ, Thøstesen LM, Kofoed P-E, et al. Effects of Bacillus Calmette-Guérin (BCG) vaccination at birth on T and B lymphocyte subsets: Results from a clinical randomized trial. Sci Rep 2017;7:12398. https://doi.org/10.1038/s41598-017-11601-6.

[26] Fink AL, Engle K, Ursin RL, Tang W-Y, Klein SL. Biological sex affects vaccine efficacy and protection against influenza in mice. Proc Natl Acad Sci U S A 2018;115:12477–82. https://doi.org/10.1073/pnas.1805268115.

[27] Márquez EJ, Chung C-H, Marches R, Rossi RJ, Nehar-Belaid D, Eroglu A, et al. Sexual-dimorphism in human immune system aging. Nat Commun 2020;11:751. https://doi.org/10.1038/s41467-020-14396-9.

[28] Porta C, Rimoldi M, Raes G, Brys L, Ghezzi P, Di Liberto D, et al. Tolerance and M2 (alternative) macrophage polarization are related processes orchestrated by p50 nuclear factor kappaB. Proc Natl Acad Sci U S A 2009;106:14978–83. https://doi.org/10.1073/pnas.0809784106.

[29] Lardone RD, Chan AA, Lee AF, Foshag LJ, Faries MB, Sieling PA, et al. Mycobacterium bovis Bacillus Calmette-Guérin Alters Melanoma Microenvironment Favoring Antitumor T Cell Responses and Improving M2 Macrophage Function. Front Immunol 2017;8:965. https://doi.org/10.3389/fimmu.2017.00965.

[30] Wieland A, Patel MR, Cardenas MA, Eberhardt CS, Hudson WH, Obeng RC, et al. Defining HPV-specific B cell responses in patients with head and neck cancer. Nature 2020. https://doi.org/10.1038/s41586-020-2931-3.

[31] Kroeger DR, Milne K, Nelson BH. Tumor-Infiltrating Plasma Cells Are Associated with Tertiary Lymphoid Structures, Cytolytic T-Cell Responses, and Superior Prognosis in Ovarian Cancer. Clin Cancer Res an Off J Am Assoc Cancer Res 2016;22:3005–15. https://doi.org/10.1158/1078-0432.CCR-15-2762.

[32] Carè A, Bellenghi M, Matarrese P, Gabriele L, Salvioli S, Malorni W. Sex disparity in cancer: roles of microRNAs and related functional players. Cell Death Differ 2018;25:477–85. https://doi.org/10.1038/s41418-017-0051-x.

[33] Shen Z, Rodriguez-Garcia M, Patel M V, Barr FD, Wira CR. Menopausal status influences the expression of programmed death (PD)-1 and its ligand PD-L1 on immune cells from the human female reproductive tract. Am J Reprod Immunol 2016;76:118–25. https://doi.org/10.1111/aji.12532.

[34] Kates M, Matoso A, Choi W, Baras AS, Daniels MJ, Lombardo K, et al. Adaptive Immune Resistance to Intravesical BCG in Non-Muscle Invasive Bladder Cancer: Implications for Prospective BCG-Unresponsive Trials. Clin Cancer Res an Off J Am Assoc Cancer Res 2020;26:882–91. https://doi.org/10.1158/1078-0432.CCR-19-1920.

[35] DeNardo DG, Ruffell B. Macrophages as regulators of tumour immunity and immunotherapy. Nat Rev Immunol 2019;19:369–82. https://doi.org/10.1038/s41577-019-0127-6.

[36] Humphrey PA, Moch H, Cubilla AL, Ulbright TM, Reuter VE. The 2016 WHO Classification of Tumours of the Urinary System and Male Genital Organs-Part B: Prostate and Bladder Tumours. Eur Urol 2016;70:106–19. https://doi.org/10.1016/j.eururo.2016.02.028.

[37] Pocock SJ, Clayton TC, Altman DG. Survival plots of time-to-event outcomes in clinical trials: good practice and pitfalls. Lancet (London, England) 2002;359:1686–9. https://doi.org/10.1016/S0140-6736(02)08594-X.

