## Supplemental figures and tables for "Sexual dimorphism in outcomes of non-muscle invasive bladder cancer: a role of CD163+ M2 macrophages, B cells and PD-L1 immune checkpoint"

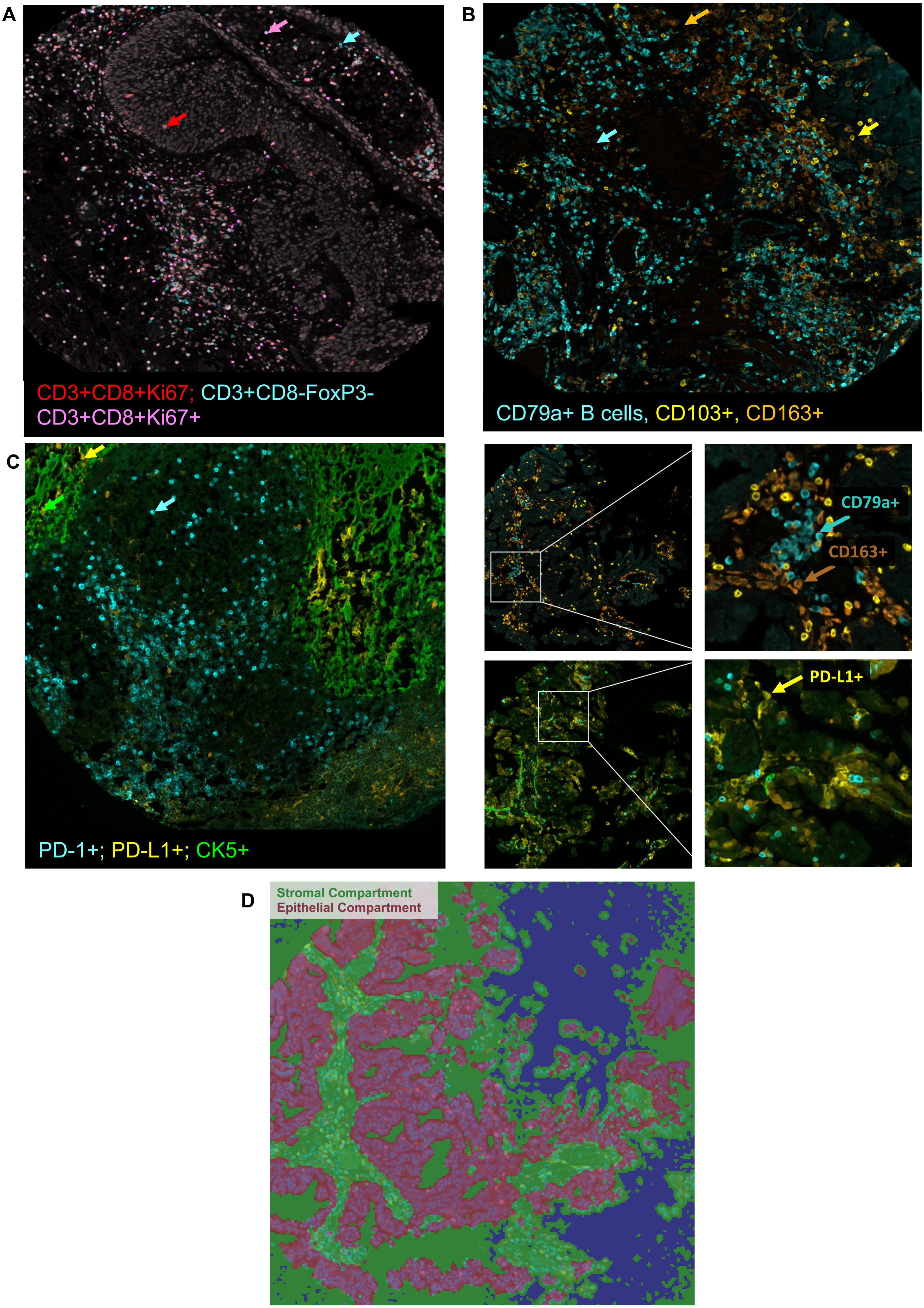


**Supplementary Figure 1. Multiplex immunofluorescence staining and automated tissue segmentation of representative tumor core.**

2A

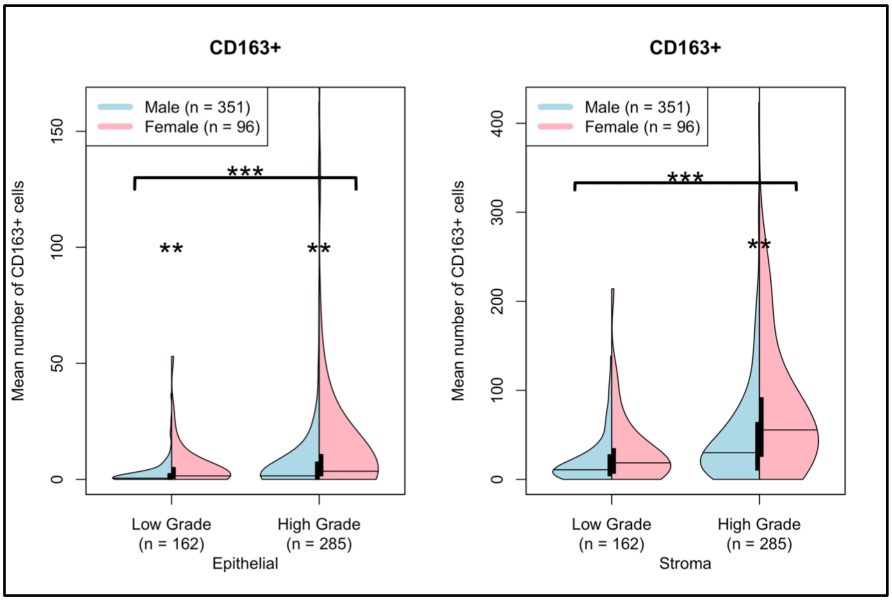


2B

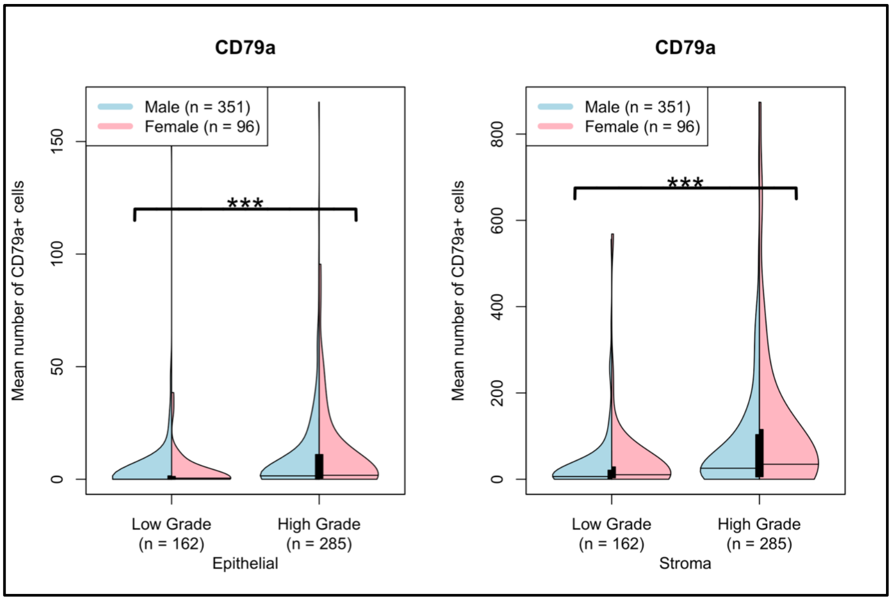


2C

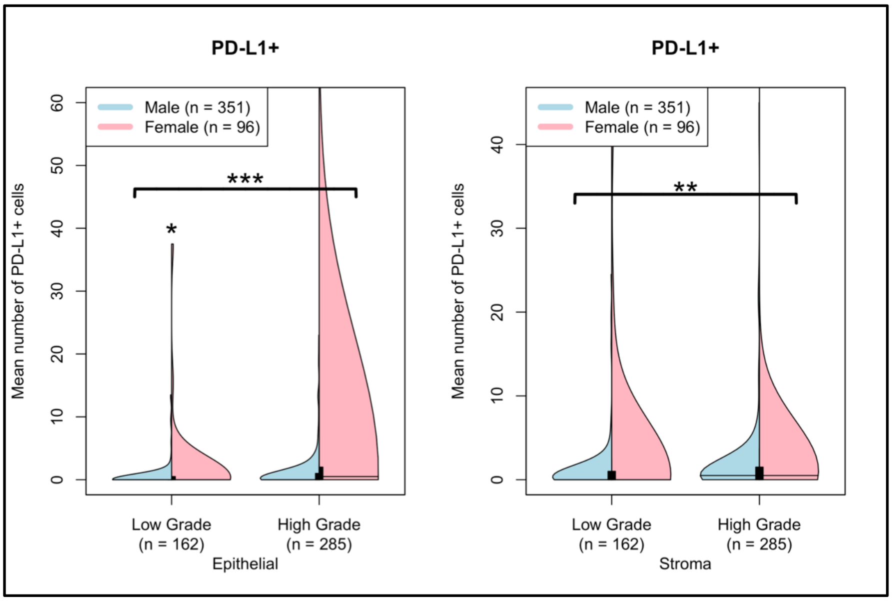


**Supplementary Figure 2: Profiles of CD163, CD79a and PD-L1 in tumors from BCG naïve patients**

3A

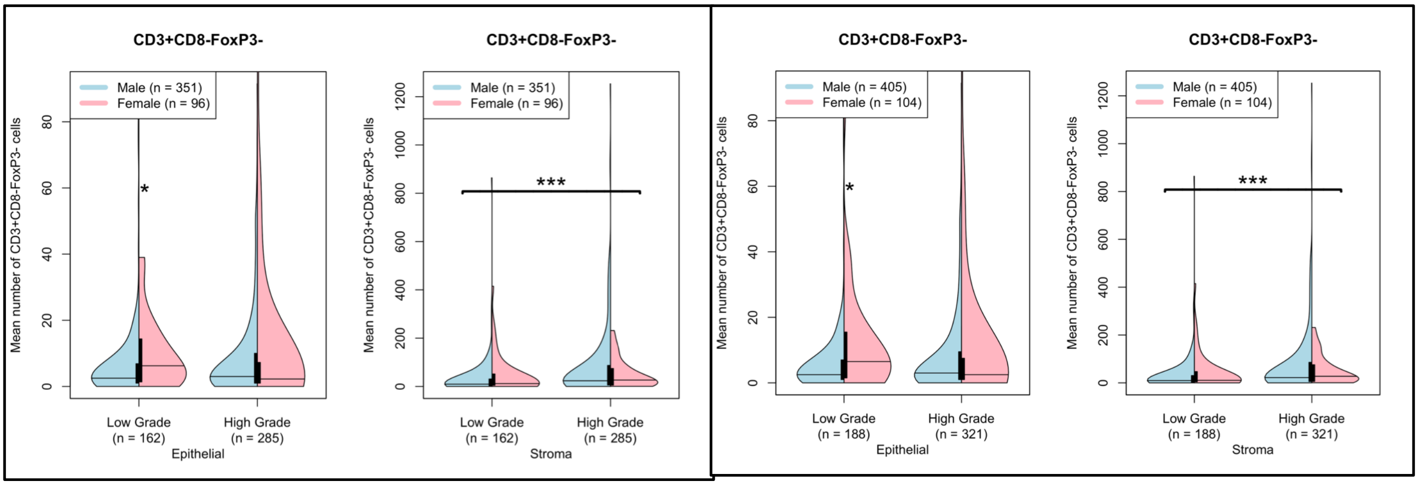


BCG naïve tumors

Overall cohort

3B

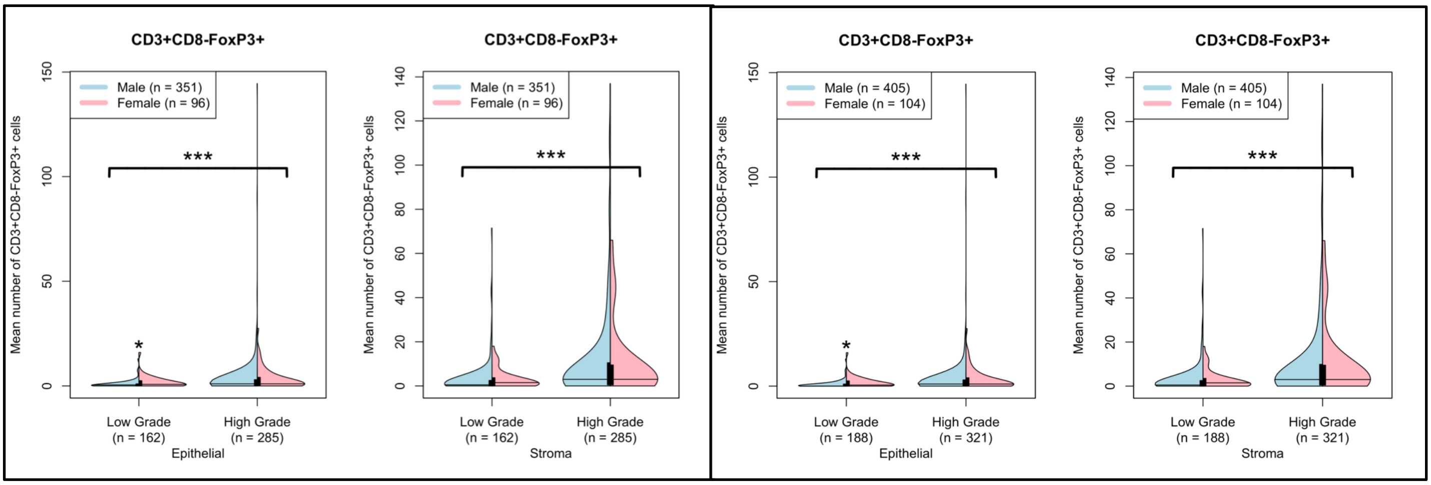


BCG naïve tumors

Overall cohort

**Supplementary Figure 3: Profiles of T helper (3A) and T regulatory (3B) cells in tumors from BCG naïve patients and overall cohort**

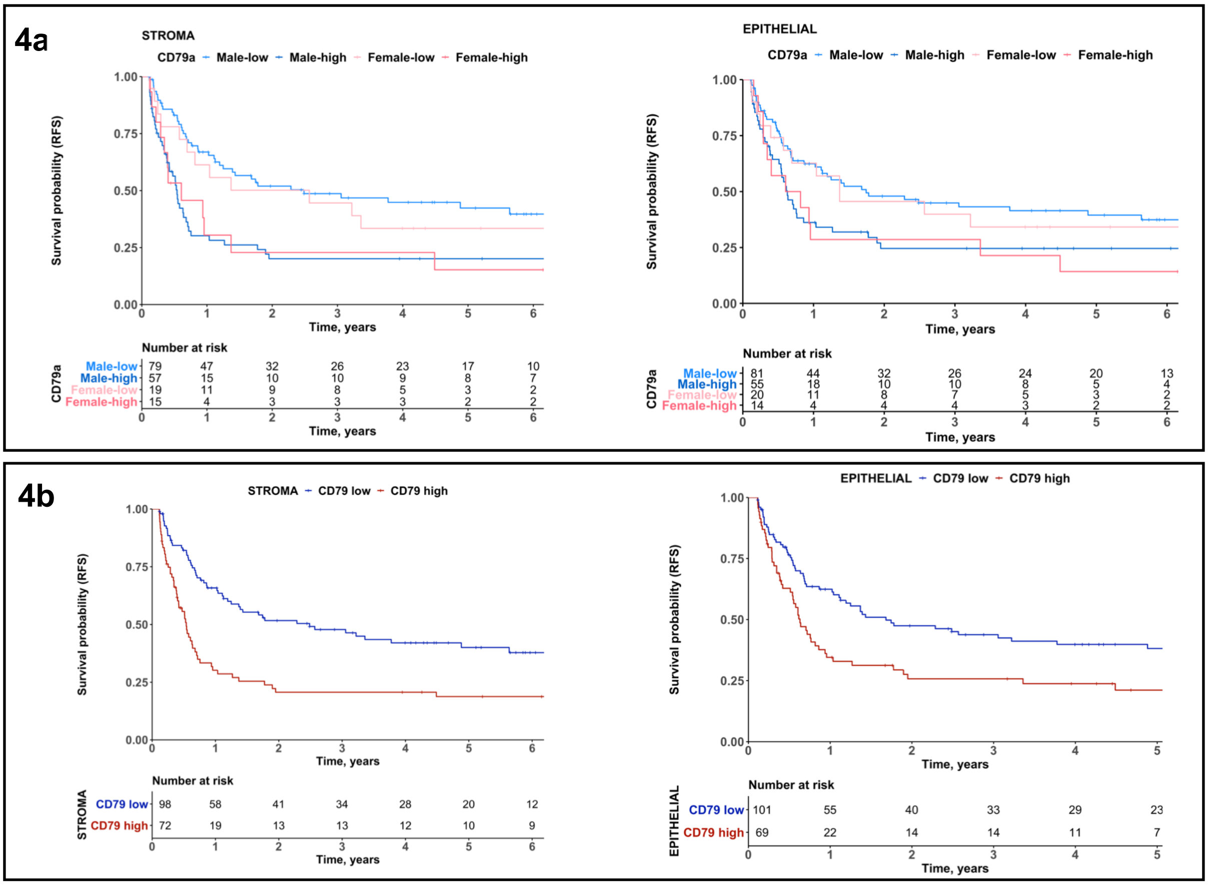


**Supplementary Figure 4. CD79a+ B cell density is associated with recurrence free survival in patients with high-grade NMIBC**

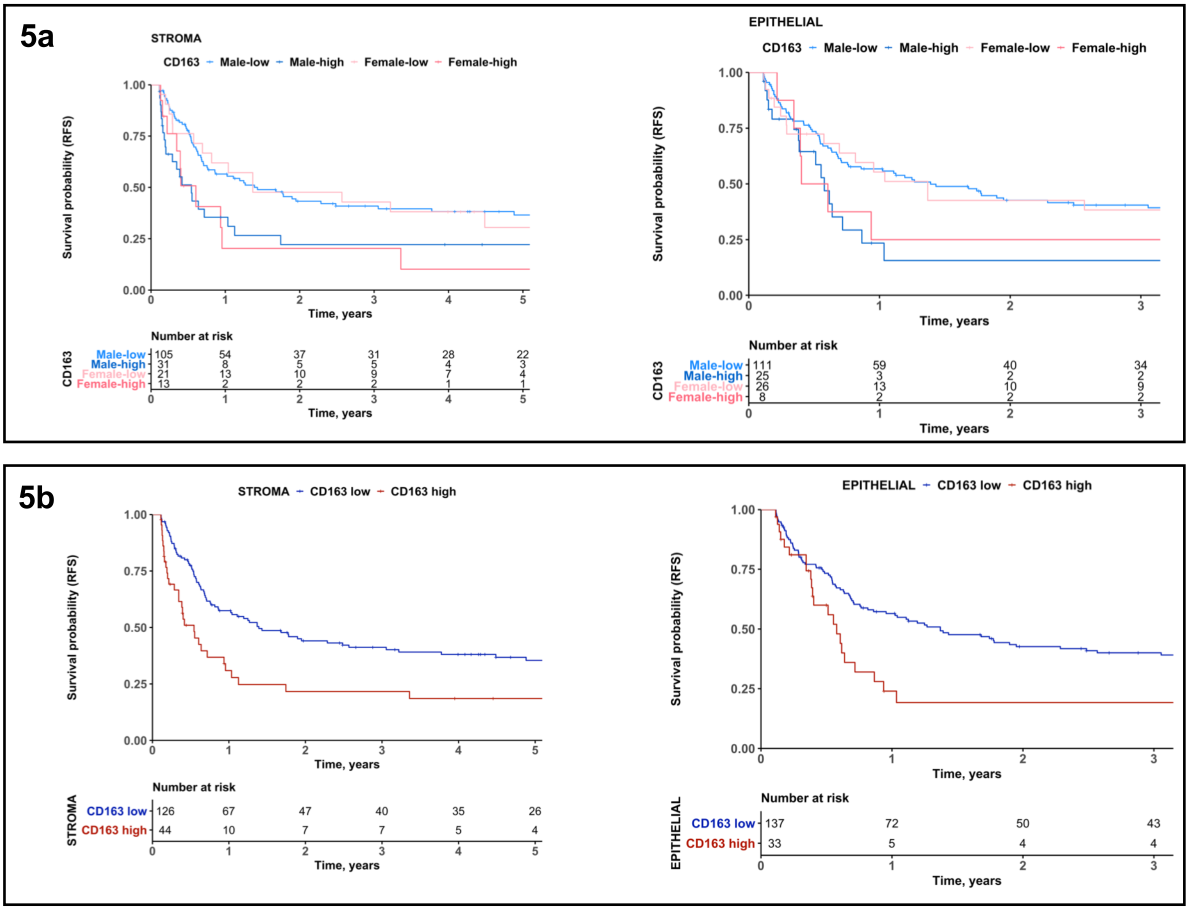


**Supplementary Figure 5. Higher CD163+ cell infiltration is associated with shorter recurrence free survival in patients with high-grade NMIBC**

**Table S1: Patient characteristics for cohort (n = 332)**

| | | |
|---|---|---|
| **Age at Diagnosis** | Mean | 71.18 |
| | Median | 72 |
| | Range | 34-94 |
| **Sex** | Male | 259 |
| | Female | 73 |
| **Initial Diagnosis, Stage and Grade** | Ta, Low Grade | 134 |
| | Ta, High Grade | 120 |
| | T1, High Grade | 78 |
| **Previous BCG** | Yes | 25 |
| | No | 292 |
| | Unknown | 15 |
| **BCG** | Yes | 180 |
| | No | 137 |
| | Unknown | 15 |
| **AUA Risk Score** | Low | 94 |
| | Intermediate | 72 |
| | High | 166 |
| **Progression to T2** | Yes | 36 |
| | No | 273 |
| | Unknown | 23 |
| **Recurrence** | < 1 year | 123 |
| | >1 year | 72 |
| | Never | 137 |
| **Median Follow-up** | | 56.2 months (4.7 years) |

**Table S2: Sample characteristics for cohort (n = 509)**

| | | |
|---|---|---|
| **Stage and Grade** | Ta, Low Grade | 188 |
| | Ta, High Grade | 165 |
| | T1, High Grade | 156 |
| **Sex** | Male | 405 |
| | Female | 104 |

**Table S3**: Optimized log-rank thresholds for recurrence-free survival for individual immune markers

| | **All high-grade patients** | **High grade patients recurring within one year** |
|---|---|---|
| **CD163-Stroma** | 64 | 65 |
| **CD163-Epithelial** | 13 | N.S |
| **CD79a-Stroma** | 35 | N.S |
| **CD79a-Epithelial** | 3 | N.S |

N.S- No significant differences in recurrence-free survival found for any cut-of
